# A UAV-based high-throughput phenotyping approach to assess time-series nitrogen responses and identify traits associated genetic components in maize

**DOI:** 10.1101/2021.05.24.445447

**Authors:** Eric Rodene, Gen Xu, Semra Palali Delen, Christine Smith, Yufeng Ge, James Schnable, Jinliang Yang

**Affiliations:** Department of Agronomy and Horticulture, University of Nebraska-Lincoln, Lincoln, NE 68583, USA; Center for Plant Science Innovation, University of Nebraska-Lincoln, Lincoln, NE 68583, USA; Department of Biological Systems Engineering, University of Nebraska-Lincoln, Lincoln, NE 68583, USA

## Abstract

Advancements in the use of genome-wide markers have provided new opportunities for dissecting the genetic components that control phenotypic trait variation. However, cost-effectively characterizing agronomically important phenotypic traits on a large scale remains a bottleneck. Unmanned aerial vehicle (UAV)-based high-throughput phenotyping has recently become a prominent method, as it allows large numbers of plants to be analyzed in a time-series manner. In this experiment, 233 inbred lines from the maize diversity panel were grown in a replicated incomplete block under both nitrogen-limited conditions and following conventional agronomic practices. UAV images were collected during different plant developmental stages throughout the growing season. A pipeline for extracting plot-level images, filtering images to remove non-foliage elements, and calculating canopy coverage and greenness ratings based on vegetation indices (VIs) was developed. After applying the pipeline, about half a million plot-level image clips were obtained for 12 different time points. High correlations were detected between VIs and ground truth physiological and yield-related traits collected from the same plots, i.e., Vegetative Index (VEG) vs. leaf nitrogen levels (Pearson correlation coefficient, *R* = 0.73), Woebbecke index vs. leaf area (*R* = -0.52), and Visible Atmospherically Resistant Index (VARI) vs. 20 kernel weight – a yield component trait (*R* = 0.40). The genome-wide association study was performed using canopy coverage and each of the VIs at each date, resulting in *N* = 29 unique genomic regions associated with image extracted traits from three or more of the 12 total time points. A candidate gene *Zm00001d031997*, a maize homolog of the *Arabidopsis HCF244* (*high chlorophyll fluorescence 244*), located underneath the leading SNPs of the canopy coverage associated signals that were repeatedly detected under both nitrogen conditions. The plot-level time-series phenotypic data and the trait-associated genes provide great opportunities to advance plant science and to facilitate plant breeding.

## Introduction

Crop improvement has long been an important goal in agriculture and much research has been conducted towards improving plant traits such as grain yield and nitrogen use efficiency. High-throughput phenotyping is an important and recent development for agriculture, it enables the quick and economical scoring of crop phenotypes on a large scale, potentially accelerating efforts towards further crop improvement via plant breeding. Numerous data acquisition methods have been employed to collect data on crop plants for high throughput phenotyping including the use of RGB, thermal, infrared, and hyperspectral cameras carried on either unmanned aerial vehicles (UAVs) or orbital platforms^[1,2,3,4]^. UAV imaging can collect data from all plots in large field experiments more frequently than a person can walk the field and manually inspect plant plots. Aerial images can also identify differences in plot-level plant health which may not be readily apparent from visual inspection on the ground. Furthermore, different crop genotypes often may react differently to the present growing conditions.

UAV imagery has seen extensive use in agriculture for a variety of purposes. It has been investigated for estimating plant height^[5,6,7,8]^ and detecting genetic loci influencing this trait^[6]^. Numerous papers have investigated using UAVs to collect spectral data from crop field trials, which can be used to predict yield^[9,10]^, generate crop surface models^[9]^, estimate plant height^[7,8]^ or biomass^[8]^, and detect nitrogen stress^[11]^. Spectral data is often assessed in the form of vegetation indices (VIs).

Numerous VIs have been defined and evaluated for quantifying different plant properties from the sensor data collected from cameras carried by aerial or orbital platforms^[1,2,3]^. For cameras which incorporate measurement of near-infrared or red edge wavelengths, these include the Normalized-Difference Vegetation Index (NDVI)^[12]^, the Soil-Adjusted Vegetation Index (SAVI)^[13]^, and the Leaf Water Content Index (LWCI)^[14]^. Indices also exist for use with RGB image data. As a result, aerial photography presents a simple way to rate the environmental response of each genotype through the plants’ greenness ratings derived from such RGB-based indices^[1,2,15]^. The greenness ratings can also correlate with the onset of flowering or senescence^[1,2]^. This provides a high-throughput method by which various genotypes can easily be rated, and with proper understanding of these correlations, the developmental progression of plants in individual plots can be predicted based on the photography. Several previous studies have also investigated the feasibility of employing VIs to predict crop yield^[16,17,18,19,20,21]^. Various different indices have been developed, each one with unique properties and applications. NDVI was found to be generally more effective than the Visible Atmospherically Resistant Index (VARI) and the Triangular Greenness Index (TGI) in assessing field health although there were individual use cases where different indices were more effective^[22]^. Indices with strong correlations to leaf area index (LAI), such as the VARI and NDVI, were found to be the most effective at predicting grain yield in rice, but indices incorporating near-infrared or red edge reflectance also tended to perform better than RGB-based indices^[23]^.

Before the image ratings can be calculated, it was suggested to eliminate debris, shadows, and bare soil^[24]^ – all features unrelated to the crop foliage, and which will therefore affect the accuracy of the greenness ratings. A common method to eliminate unwanted pixels in crop images is to use VIs. Probably the most commonly-used index is the NDVI^[12]^, however, this method relies on near-infrared spectral data, which provided challenges for cost-effectively accessible RGB data.

In this study, maize RGB imagery data was collected for a maize diversity panel using UAV-based high-throughput phenotyping from a replicated field experiment with standard and nitrogen deficient management practices. A pipeline was developed using commercial software to extract images pertaining to individual genotypes in each plot. With these plot-level images, several RGB-based VI calculations were developed to compare nitrogen responses for different genotypes and over the different plant developmental stages. Correlation studies of the VIs with the plant physiological and yield-related traits suggested some VIs can be employed as indicators for predicting the phenotypic traits of interest. Finally, genome-wide association study (GWAS) was conducted to identify genetic loci associated with image extracted traits. This plot-level high throughput phenotyping method and the identified genetic loci have the great potential to benefit plant breeding.

## Results

### A computational pipeline to extract plot-level images from a replicated field experiment

In the summer of 2019, a maize diversity panel consisting of 233 inbred lines drawn from the Buckler-Goodman Association Panel^[25]^ was grown at a UNL experimental station based on an incomplete block design with two replications (**Figure 1A**). Each replication included two main plots either with or without nitrogen (N) fertilizer treatment. For each plot, four split plots were blocked by plant height and maturity (see **Materials and Methods**). Each split plot was further subdivided into three split plot blocks. A hybrid check was randomized within each split plot block as a subplot. During the growing season, a Phantom 4 Pro UAV equipped with a red, green, and blue (RGB) sensor was flown at about 22-27 m above ground and captured between 210 and 360 partially overlapping images each flight. A total of 12 flights were conducted between July 6 and Sept 5. An average of approximately 2,500 clipped plot-level RGB images per quadrant per date were generated from the initial UAV imagery. The original UAV images taken for this study are available at CyVerse (DOI: 10.25739/4t1v-ab64).

**Figure 1:**
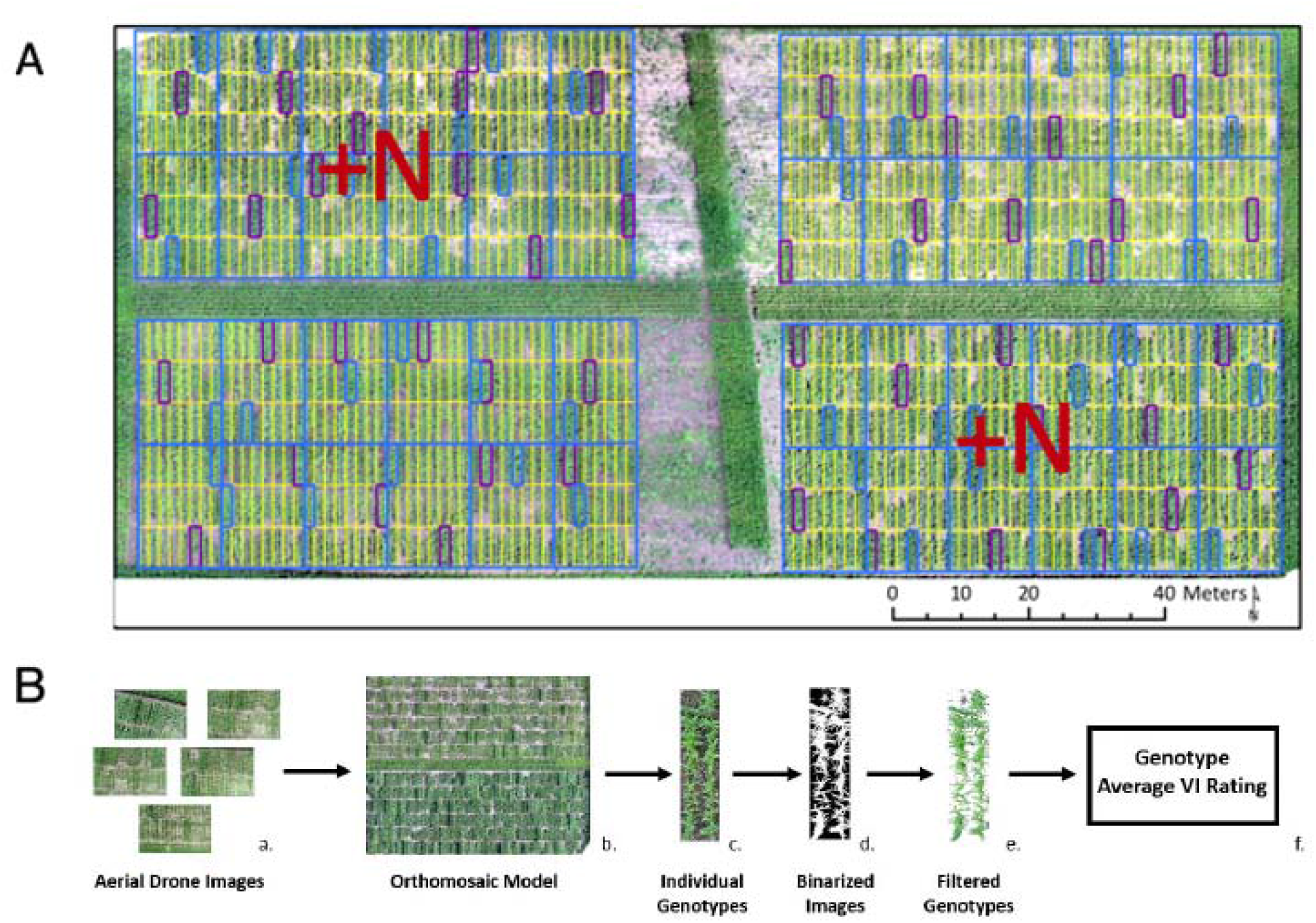
Field design and image data processing pipeline. (**A**) The orthomosaic map created from the UAV-based imagery. Yellow grid lines indicate the positions of individual two-row plots. Purple and blue rectangles indicate the control plots. Inorganic nitrogen fertilizer (+N) was applied to the top left and bottom right quadrants of the field prior to planting. (**B**) Workflow diagram of the image data processing. Individual UAV images (a) were used to generate an orthomosaic model (b). The partitioned genotypes (c) were extracted from the original UAV images, generating multiple replicates per plot. These images were then filtered using binary masks (d) to remove non-foliage pixels (e). The resulting images were then processed using a variety of vegetation indices to calculate average greenness ratings for each genotype at the point in the growing season the images were collected (f).

In order to obtain phenotypic values for each genotype (i.e., the two-row subplots) from these images, a computational pipeline was developed to process the UAV data into plot-level images (see **Materials and Methods**). Briefly, 1) the aerial UAV images collected from each day were used to form an orthomosaic model of the entire field; 2) clipped images of individual subplots, depicting separate genotypes, were extracted using the commercial software Plot Phenix^[26]^; 3) these plot-level images were filtered to remove non-foliage pixels; 4) the resulting binary mask is then used to avoid non-foliage pixels when calculating phenotypic values, i.e., VIs (**Figure 1B**). After the individual plot images are extracted, a pixel filtration procedure was implemented to remove non-foliage pixels (see **Materials and Methods**), as including non-foliage pixels in these calculations will affect accuracy of the greenness ratings ^[24]^.

### Elevated Canopy Coverage in the Nitrogen Applied Field

Each designated two-row plot was captured in multiple UAV images taken in different positions above the field. The viewing angle between replicated photos taken of a single plot during the same flight is highly variable significantly altering estimates of canopy coverage based on the binary masks (as seen in **Supplementary Figure S1)**. The most nadir image in the set of images collected for each individual plot was selected as the reference replicate, and these reference images were used to calculate percent canopy coverage for each of the two row plots at each time point.

The canopy coverages of the hybrid check in the +N quadrants were higher than the -N quadrants, although there is overlap in the overall ranges of values (**Figure 2A)**. The differences were statistically significant in earlier days, i.e., July 6 (8.0% difference, *p*-value = 9.6e-11) and Aug 12 (10% difference, *p*-value = 7.9e-32), but the differences became insignificant as plants reached maturity and began to senescence, i.e., Sept 5 (1.6% difference, *p*-value = 0.06). This reduced N response was expected since the N fertilizer was applied before planting, the hybrid check would reduce its positive reaction to nitrogen treatment as they develop. Similarly, the pattern of reduced N reactions was observed for the *N* = 233 diverse inbred lines (**Figure 2B**). In July 6, the average ratio of canopy coverage (+N/-N) was 1.2 (paired t test, *p*-value = 1.7e-37). The ratios reduced to 1.1 in the August 12 (paired t test, *p*-value = 3.7e-05) and stayed around this value later on and become insignificant in Aug 14 and Sept 5th (**Figure 2B**). Although the results suggested most of the genotypes react positively in terms of canopy coverage under +N treatment, i.e., 17 genotypes exhibiting a ratio > 1.5, there were a number of genotypes that reacted negatively to nitrogen, i.e., 76 genotypes showing a ratio < 0.8 (**Supplementary Figure S2A**). The responses of these genotypes to N treatments seemed largely consistent throughout the growing season (**Supplementary Figure S2B**).

**Figure 2:**
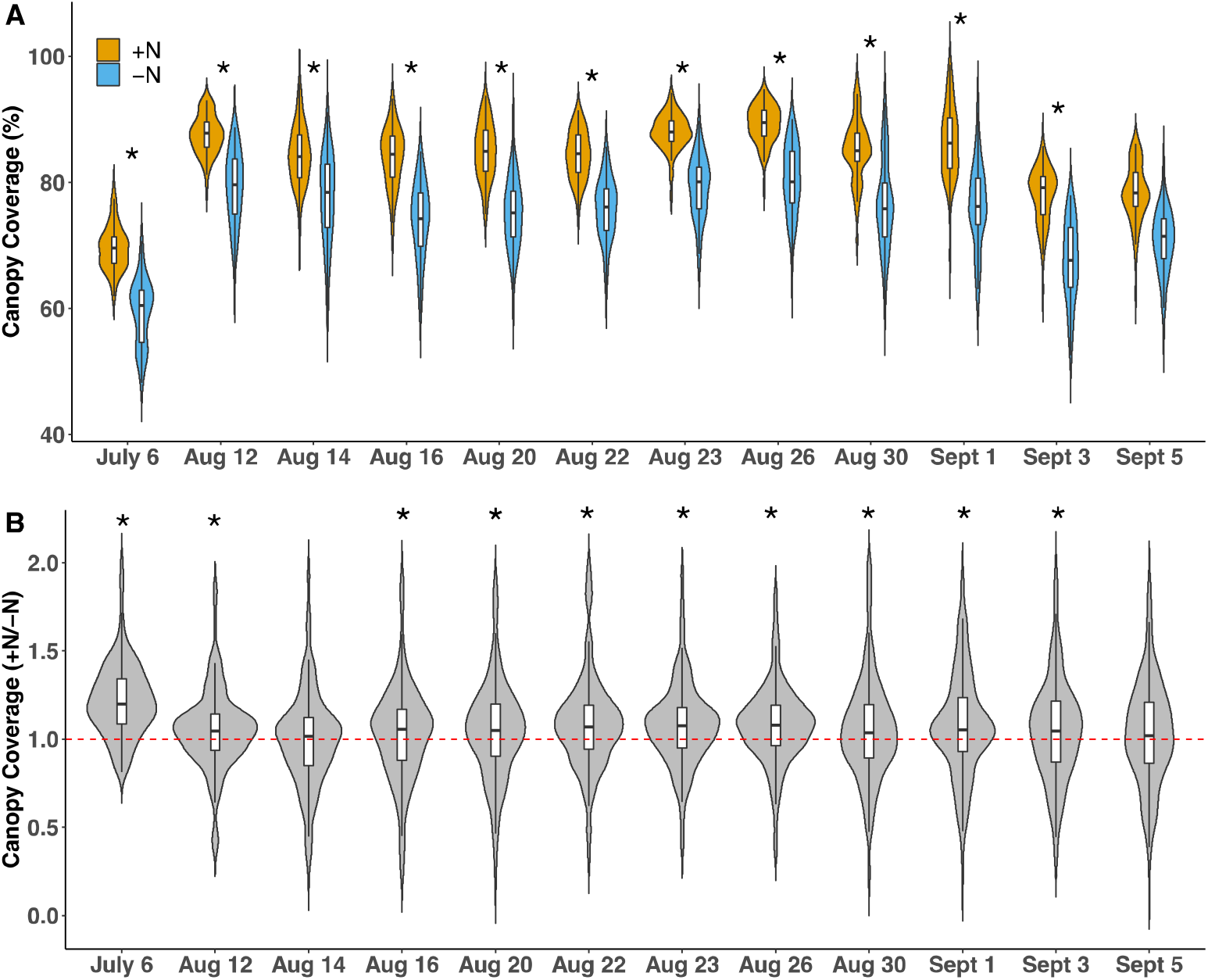
Time series canopy coverages throughout the plant growing season. (**A**) Canopy coverage for the hybrid checks with (+N) and without (-N) nitrogen treatment. The reference replicate images of each of the plots containing the hybrid check were used on each date. (**B**) The ratio of canopy coverage (+N/-N) for the *N* = 233 maize genotypes throughout the season. Asterisk indicates *p*-value < 0.05.

### Vegetation indices (VIs) correlated with leaf nutrient traits and ear-related traits

Eight VIs were calculated from the images collected in this study: the Excess Green Index (ExG), the Red-Green-Blue Vegetation Index (RGBVI), the Normalized Green-Red Difference Index (NGRDI), the Green Leaf Index (GLI), the Modified Green-Red Vegetation Index (MGRVI), the Visible Atmospherically Resistant Index (VARI), the Vegetative Index (VEG), and the Woebbecke Index (**Table 1**). Consistent with previous reports^[27]^, the VIs were sensitive to the light value and color saturation of the images (**Supplementary Figure S3**). After dividing by light and color saturation values, the normalized VIs can clearly distinguish the +N and -N quadrants using the hybrid check, where most of the VIs (7/8) exhibited higher values under +N treatment and the Woebbecke index showed significantly lower values (**Supplementary Figure S4**). Consistently, average ratios of the VIs (+N/-N) for the 233 genotypes were deviated from 1 (**Supplementary Figure S5**), suggesting their sensitivity in detecting N treatments.

**Table 1:**
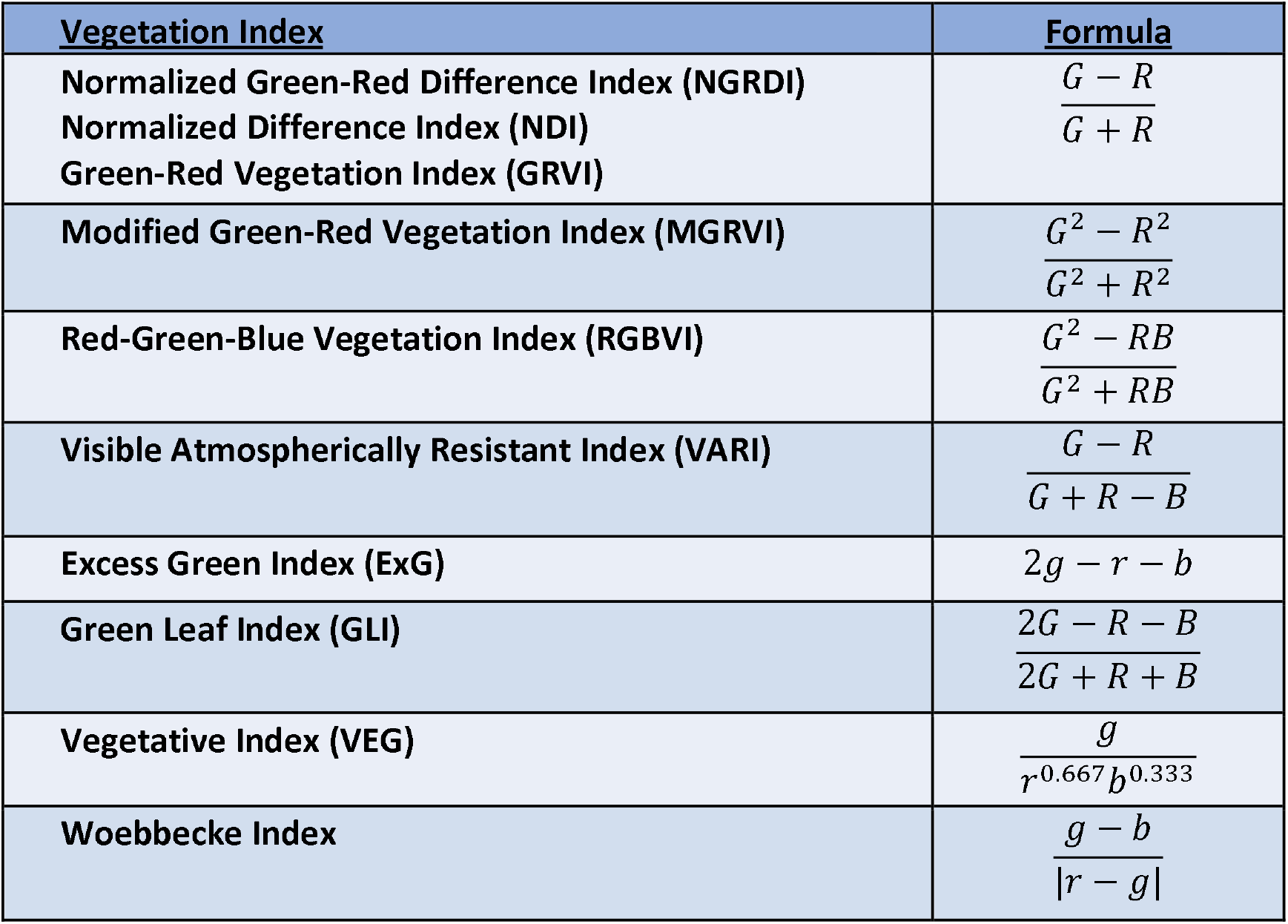
A listing of the vegetation indices used in this study, with their formulas. The variables *R, G*, and *B* represent the values of the red, green, and blue channels, respectively, of a given pixel. While the variables *r, g*, and *b* represent these values normalized by dividing the red, green, or blue channel of a given pixel by the sum of the values of all three channels (see **Supplementary Information** for more details).

The VI values were correlated with measurements of leaf nitrogen levels, leaf areas, and 20 kernel weights from sampled leaves and mature ears collected from the same field to evaluate the reliability of using VIs to predict physiological and yield-related traits (see **Materials and Methods**). Overall, statistically significant correlations were observed with mean Pearson correlation coefficients of *R* = 0.27±0.17 for leaf nitrogen level, *R* = 0.24±0.12 for leaf areas, and *R* = 0.16±0.08 for 20 kernel weight (**Figure 3A**). Over the growing season, the degree of correlation between VIs and ground truth traits exhibited remarkable fluctuations. The coefficient values peaked on Aug 12 and Aug 22 and dramatically reduced on Aug 16 and Aug 26, partly because images were taken late in these days (between 7:30 and 8:00 pm, local time), which had a noticeable effect on the light values of the images. Aug 12 had light values between 9.97 and 10.97, Aug 22 had light values between 10.64 and 11.95, but Aug 16 and 26 both had light values between 6.97 and 8.32. Notably, unlike the other VIs, the VEG performed better on these unusual days, especially at predicting leaf area.

**Figure 3:**
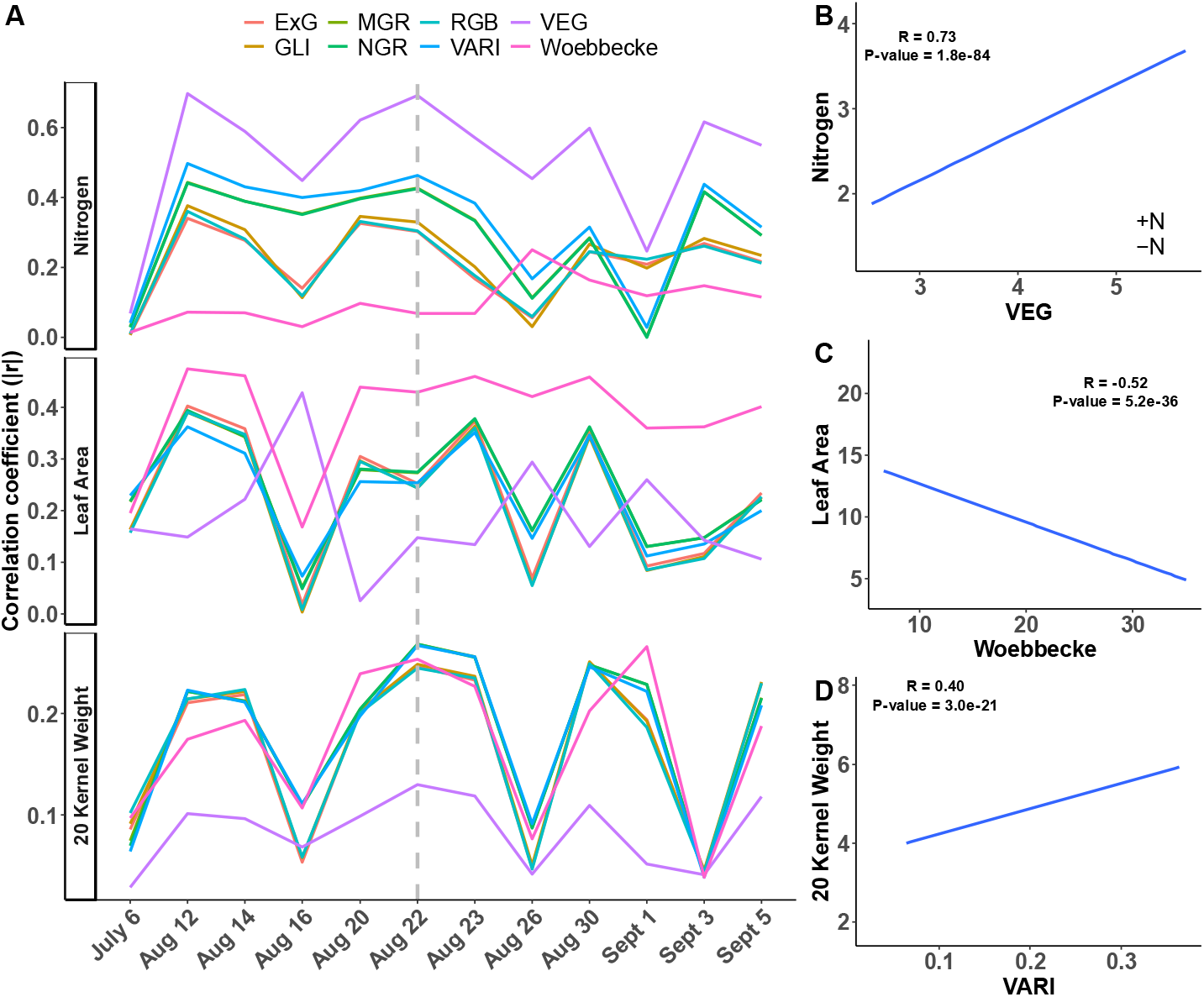
Correlation of the VIs with physiological and yield-related traits. (**A**) Time-series correlation coefficients of the eight VIs with leaf nitrogen levels, leaf areas, and 20 kernel weights. Grey dashed line indicates the selected date for further investigation. The initials MGR, RGB, and NGR refer to the MGRVI, RGBVI, and NGRDI, respectively. Scatter plots of the VEG vs. nitrogen (**B**), Woebbecke vs. leaf area (**C**), and 20 VARI vs. kernel weight (**D**) in the selected date of August 22. Blue lines denote the fitted regression line using a linear model and grey shaded areas indicate the 95% confidence intervals. Text in the plots show the Pearson correlation coefficients and associated *p*-values.

The eight VIs varied substantially in their predictabilities for different traits. The VEG outperformed other indices for the leaf nitrogen level, exhibiting the highest *R* = 0.73 on Aug 22 (**Figure 3B**). The Woebbecke index, although displaying a negative correlation with the traits, the absolute *R* values of Woebbecke were mostly above other indices for predicting the leaf area trait (**Figure 3C**). It became more challenging to predict the yield-related trait, i.e., 20 kernel weight. Occasionally, the VARI exhibited a slightly higher correlation efficient, i.e., in Aug 22 (**Figure 3D**). But the overall performance for yield-related trait prediction was considerably lower than the leaf nitrogen level and leaf area traits.

### Genome-wide association studies (GWAS) identified canopy coverage and VI-associated loci

GWAS was conducted using a mixed linear model for the time-series canopy coverage and VI traits and a set of more than 20 million SNPs obtained from the whole-genome sequencing data^[28]^. Using a modified Bonferroni threshold of 1.2×10^−6^ (1/*n, n* = 769,690 is the number of independent SNPs)^[29]^ and more than five significant SNPs within a 100-kb window as a criteria (see **Materials and Methods**), a total of 14 unique regions were detected with significant associations with variation in canopy coverage (**Figure 4**) and 140 regions exhibited significant associations with VI traits (see **Supplementary Table S1** for trait-associated SNPs). For canopy coverage traits, three overlapping GWAS regions were detected between +N and -N fields, while nine (+N) and two (-N) treatment specific GWAS regions were found, suggesting genetic control for canopy coverage are moderately distinct under different N treatments. Comparing to the canopy coverage traits, fewer consistent GWAS peaks at different dates were detected for each VI trait. For example, 9 (+N) and 11 (-N) unique GWAS regions were detected for the VEG, the index exhibiting the highest correlation with leaf N content, however, very few of these regions were repeatedly detected on more than two different dates (**Supplementary Figure S6**).

**Figure 4:**
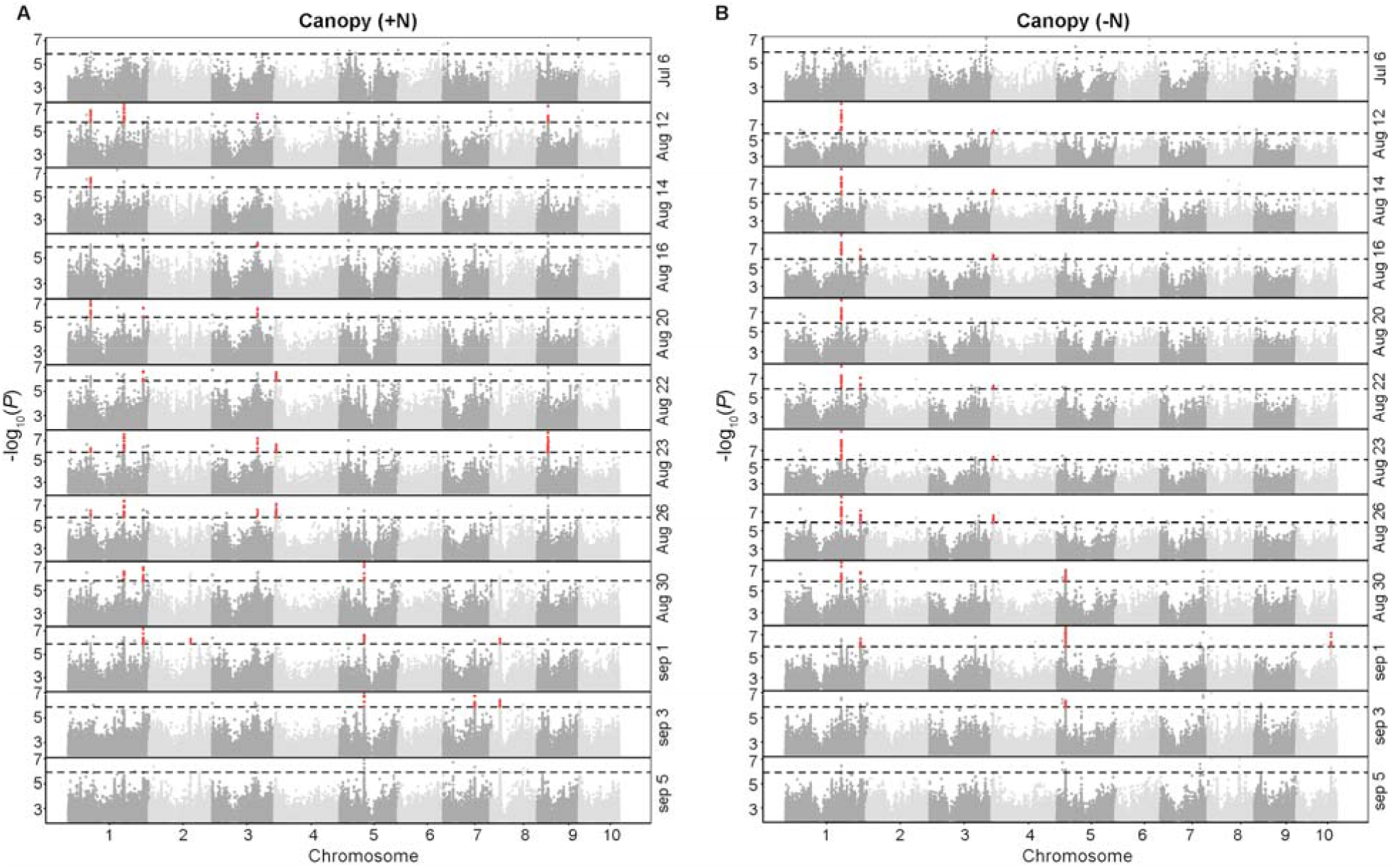
GWAS results for canopy coverage at different dates with (A) and without nitrogen (B) treatments. Red dots highlight the GWAS regions with more than five significant SNPs within a 100-kb window. Horizontal dashed lines indicate the GWAS thresholds.

A set of 29 unique regions were associated with the same image extracted feature in distinct analyses conducted on data from three or more of the twelve total time points phenotyped. Among these GWAS signals, 13 (44.8%) regions were also associated with variation in two or more distinct image extracted features (**Figure 5A**). Notably, a canopy coverage associated GWAS peak located on the Chr1 208.8-209.1 Mb region was repeatedly detected in multiple days under both N conditions (**Figure 4** and **Figure 5A**). The *Zm00001d031997* gene, located 4.6 kb downstream of the lead SNP (**Figure 5B**), is a homolog of the *Arabidopsis* gene *HCF244 (high chlorophyll fluorescence 244)* which plays a role in the assembly of photosystem II^[30]^. The abundance of mRNA transcripts derived from *Zm00001d031997* is much higher in mature leaves than in other tissues (**Figure 5C**), suggesting its potential effects on plant development.

**Figure 5:**
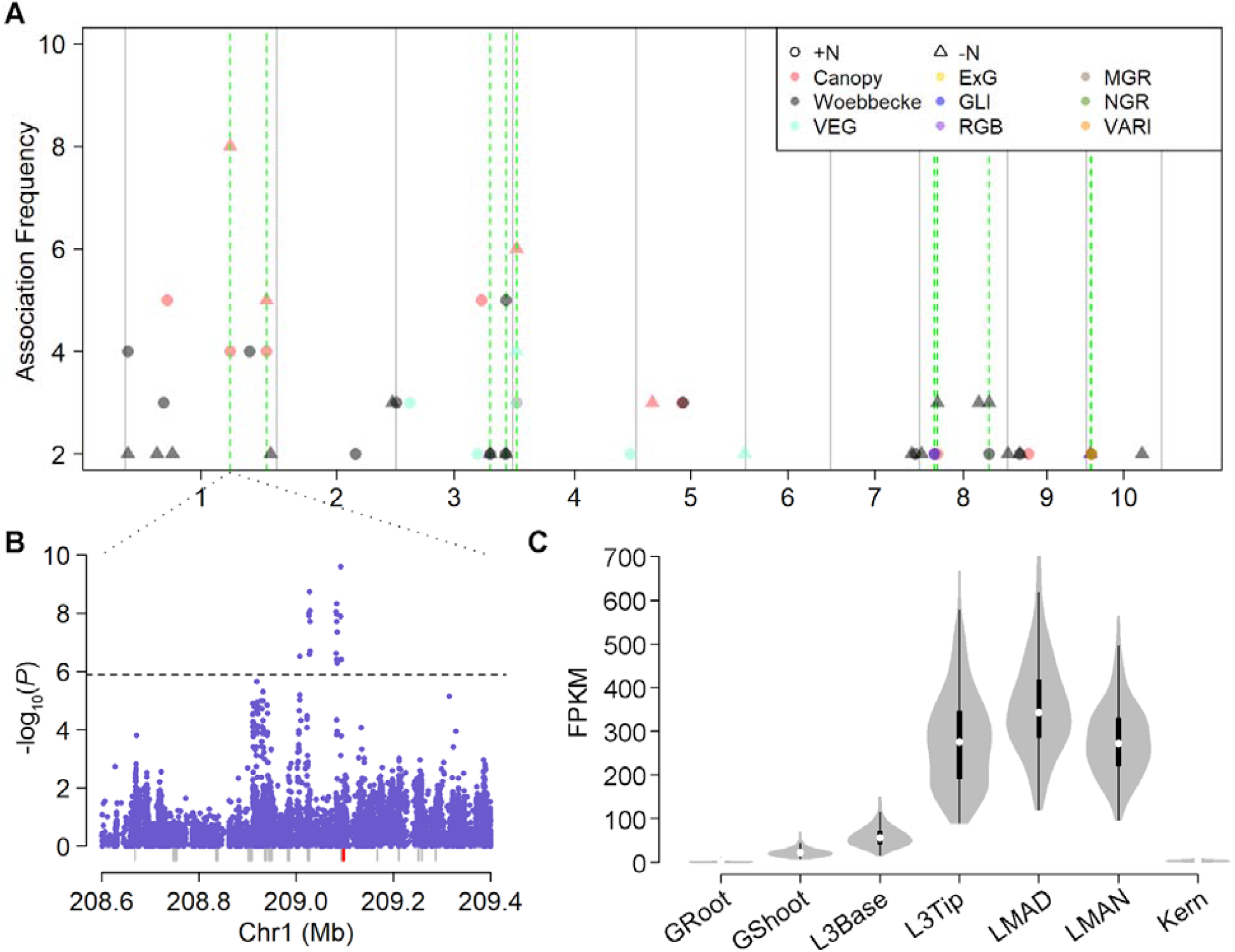
Association frequency for the canopy coverage and VI traits. (**A**) Positions and detection frequency (out of twelve total time points) of genomic intervals associated with variation in canopy coverage and the eight VI traits. Dots and triangles indicate the phenotypes collected under +N and -N conditions, respectively. The vertical green dash lines indicate regions significantly associated with multiple traits. (**B**) The zoom-in plot of the association signals at Chr1 208.8-209.1 Mb region for the canopy coverage trait collected in the -N field on August The dashed horizontal line indicates the Bonferroni-adjusted significance threshold (1.2×10^−6^). The gray rectangles indicate the gene models and the red rectangle highlights location of the candidate gene *Zm00001d031997*. (**C**) The gene expression level of *Zm00001d031997* in different tissues summarized from published data^[31,32]^.

## Discussion

In this study, a UAV-based high-throughput phenotyping pipeline has been developed to extract plot-level images, which are then used for GWAS to map a number of genetic loci associated with image-extracted traits. For the plot-level image processing pipeline, an algorithm is applied to the clipped images to filter out non-foliage elements and to calculate vegetation index averages over the surviving pixels. These greenness ratings provide a “snapshot” of that plot at specific points in the season. Some of the greenness indices, i.e., the VEG, has been demonstrated to be an effective predictor (*R* = 0.7) that can be used to predict genotype response to nitrogen treatment.

By using the VIs and canopy coverage as the phenotypic traits, about 30 uniquely trait-associated loci were identified at least twice, which may reduce the false discovery rate – a major drawback of the GWAS method^[33]^. A candidate gene, HCF244 or *Zm00001d031997*, located in one of the most significant GWAS peaks for canopy coverage trait under both +N and -N conditions. The HCF244 gene encodes a putative NAD(P)H-binding protein^[34]^. Recent evidence from studies in maize and *Arabidopsis* suggests the HCF244 is part of a protein complex involved in photosystem II (PS II) assembly and repair^[30]^. Because the protein in the PS II reaction center is subject to photodamage, the HCF244-related complex may play a key role in replacing the damaged protein with nascent one. Therefore, we speculate that the natural variations in the HCF244 gene or its regulatory modulation might affect the efficiency of the PS II protein assembling or repairing and eventually lead to the phenotypic consequences that can be detected by UAV. However, additional experiments need to be carried out to further elucidate the molecular mechanism of the genetic locus.

The study of the UAV-based image data has revealed several limitations. Virtually all of the selected VIs are highly sensitive to the color saturation of the images, producing higher values for images with greater saturation. This can easily be corrected for by normalizing the calculated VI ratings with the average saturation of each image. As was seen with the difference between the July 6 images compared to the rest of the season, it is also crucial to ensure identical camera settings are used. Furthermore, as exhibited in **Figure 3A**, it is also important to ensure the images collected are taken at a similar time of day, as images taken much later in the day will have lower light values. The Pearson correlation coefficient has been shown to be sensitive to the light values of the image, and it will be important in the future to minimize variation in this regard. It should also be noted that there will naturally be some variation in the canopy coverage estimates of the plots, as this measurement is affected not only be how large the plants are, but also how well centered the plot is in the image, the ground surface area the image encompasses (which can be affected by UAV altitude), and how much intrusion there may be from leaves in adjacent rows. However, this can also be minimized by flying at a consistent altitude on each date and making use of the reference replicate images, which in most cases are the most centered and also depict a given plot from the most nadir angle.

Given that UAV-imagery data limitations are adequately compensated for with using consistent data collection techniques, it is the hope that UAV data can be used as an accurate predictor of crop response, such as nitrogen treatment, overall plant health, and final grain yield. Additionally, the promising GWAS signals detected using the image-extracted trait provide great opportunities to advance plant sciences. Ultimately, the UAV-based pipeline and advanced statistical models have the potential to greatly benefit crop improvement in the future.

## Materials & Methods

### Field Experimental Design and UAV Data Acquisition

The plants were grown at the Havelock Research Farm of the University of Nebraska-Lincoln on the east edge of Lincoln, NE. The field was partitioned into four quadrants (NE, SE, NW, and SW), with two quadrants receiving a standard field treatment of nitrogen (the +N quadrants, NE and SW), and two quadrants receiving no treatment (the -N quadrants, NW and SE). Each quadrant was divided into six ranges, with 42 two-row subplots to each range, for a total of 252 subplots per quadrant. Row spacing was 30 inches and within row plant spacing was six inches with 38 plants per row, resulting in total subplot dimensions of 5 ft × 20 ft. A total of 233 genotypes from the Buckler-Goodman Maize Association Panel were grown using an incomplete split plot block design with between 27 and 37 plots of the check genotype inserted per quadrant.

### Phenotypic data collection

After maturity, three ears per two-row plot were hand harvested and dried at 37 °C for 2 to 3 days. Ears were hand shelled and 20 undamaged and otherwise representative kernels were collected from the bulked seed and weighed in order to determine 20 kernel weight.

### UAV Imagery Data Processing

A total of 12 flights were conducted on dates between July 6 and September 5 employing a Phantom 4 Pro UAV equipped with a red, green, and blue (RGB) sensor with a resolution of 5,472 × 3,648 pixels. All raw image data is being made available via CyVerse (DOI: 10.25739/4t1v-ab64). The resulting individual images were employed to construct separate orthomosaic models of the field for each time point using Plot Phenix^[26]^. Plot boundaries were manually defined for each time point. Plot Phenix was then used to extract the cropped original images of each plot from the raw images employed to generate the original orthomosaic. A given plot is often captured multiple times from different angles as the UAV makes its way across the field. Phenix will designate one of these replicates as that plot’s reference replicate, which is the most nadir image available out of those replicates. Extracted plot images were of varying resolution (typically between 250 × 1000 and 300 × 1200 pixels). The reference image designated by Phenix for each plot was employed for downstream analyses.

### Vegetation Indices Calculation

The process used in this study is to extract clipped images representing individual plots from aerial photographs, filter the plots to eliminate soil, shadows, and debris, and calculate a series of greenness ratings for each plot to be able to track fluctuations in the ratings throughout the season (see **Supplementary Information** for more details).

### Genome-Wide Association Study (GWAS)

The genotype of the maize diversity panel was downloaded from maize HapMap3^[28]^ with AGPv4 coordinates. After filtering out SNPs with minor allele frequency ≤ 0.05 among the 231 lines phenotyped in this study, approximately 21 million SNPs were retained for further analysis. GWAS was conducted using a mixed linear model (MLM) implemented in GEMMA (v 0.98.3)^[35]^. In the GWAS model, the first three principal components calculated by PLINK 1.9^[36]^ and the kinship matrices computed by GEMMA were fitted as fixed and random effects to control for the population structure and genetic relatedness, respectively. The threshold for the significant association SNPs was set to 1.2×10^−6^ (1/*n, n* = 769,690 is the number of independent SNPs). Here, the independent SNP number was determined by using PLINK 1.9 with the *indep-pairwise* option (window size 10 kb, step size 10, *r*^2^ ≥ 0.1). GWAS peaks were then determined by considering a window 50 kb upstream and downstream of the significant SNPs. Overlapping regions were merged, and the regions with more than five significant SNPs were defined as high confidence association regions.

## Acknowledgements

This project is supported by the National Science Foundation under award number OIA-1557417 for Center for Root and Rhizobiome Innovation (CRRI). J.Y. is supported by the Agriculture and Food Research Initiative Grant number 2019-67013-29167 from the USDA National Institute of Food and Agriculture and the University of Nebraska-Lincoln Start-up fund.

