## Supplemental information for "A UAV-based high-throughput phenotyping approach to assess time-series nitrogen responses and identify traits associated genetic components in maize"

**Supplementary Notes**

Because the drone images only include visible color bands, indices like the NDVI^[1]^, which uses near-infrared reflectance, cannot be used. Alternatives must instead be found to work with the RGB bands of visible spectrum images. One such index is the Normalized Green-Red Difference Index (NGRDI)^[2]^, the formula for which is:

$$NGRDI=\frac{G - R}{G + R}$$

Where *G* is the green value for a given pixel, and *R* is the red value for the same pixel. Other similar real-color indices include the Red-Green-Blue Vegetation Index (RGBVI)^[3]^, Green Leaf Index (GLI)^[4]^, and Visible Atmospherically Resistant Index (VARI)^[5]^.

Perhaps the best index for filtering crop images which relies purely on RGB pixel values is the Excess Green Index (ExG)^[6]^, with the following formula:

$$ExG=2G-R-B$$

Where *G* and *R* represent the green and red values, respectively, for a given pixel, and *B* represents the pixel’s blue value. All such pixel values are normalized to be between 0 and 1 by dividing the raw pixel value by 255. An image can easily be filtered by applying a threshold to the resulting calculated values. Any pixels with values lower than the selected threshold will be eliminated. In this way, a set of indices can be computed for each pixel, and values below a certain threshold may be eliminated, as these lower values tend to be associated with pixels representing non-foliage features.

Note that the formula above differs from the original form of the ExG equation presented by Woebbecke^[6]^. His form was presented as:

$$ExG=2g-r-b$$

Where the values *r*, *g*, and *b* are calculated from the RGB values of each pixel as follows:

$$r=\frac{R}{R+G+B} , g=\frac{G}{R+G+B} , b=\frac{B}{R+G+B}$$

The values *R*, *G*, and *B* are here normalized between 0 and 1, as previously described. Our in-house experiments have shown that using the normalized RGB values instead of the *r*, *g*, and *b* values Woebbecke described results in more accurate image segmentation between foliage and non-foliage features, and so this is the version that has been used here. Specifically, this results in significantly improved ability to distinguish between foliage and shadows, something that the original formula using the *r*, *g*, and *b* values has difficulty in doing.

The plot-level images were filtered to remove non-foliage elements. This was accomplished using the Excess Green Index (ExG)^[6]^, rescaling to a 0 to 255 range to make use of whole numbers for determining thresholds, and with six different threshold values selected. In-house experiments have shown that thresholds from 135 to 140 tend to be best for total elimination of soil and shadows, while still retaining most of the plant foliage.

An example of the filtering process is as follows. If a given pixel has values of *R* = 127, *G* = 211, and *B* = 113, the ExG formula calls for the RGB values to be normalized between 0 and 1, which is accomplished by dividing them each by 255 (the maximum value in this RGB scale):

$$R^{*}=\frac{127}{255} , G^{*}=\frac{211}{255} , B^{*}=\frac{113}{255}$$

Using these values, the ExG formula is:

$$ExG=2G^{*}- R^{*}- B^{*}$$

In this example, this approximates to 1.655 – 0.498 – 0.443 ≈ 0.7137.  The version of the ExG we used normally ranges from -2 to 2, but to rescale it to range from 0 to 255, this formula is applied:

$$\left( \frac{ExG+2}{4} \right)\times255$$

Where *ExG* is the calculated ExG value for a given pixel. Adding 2 to the calculated value puts it on a scale from 0 to 4, and dividing by 4 normalizes it to a 0 to 1 scale. Finally, multiplying by 255 changes the scale to range from 0 to 255.  The approximate value 0.7137 in this example is transformed into 173.  The threshold we’ve used, as determined from in-house experiments, is 131. 173 is greater than the threshold, so this pixel would be saved in filtering the image. In this way, the image is binarized. If a given calculated ExG value is at least equal to the chosen threshold, that pixel is a 1, otherwise it’s a 0.

The vegetation indices were then calculated on the filtered images. If a pixel has been binarized to a 1 using the filter, its vegetation indices will be calculated and added to a running total. Otherwise, it is ignored, since the pixel was removed in the filtering step. After all of the pixels in the image are processed in this way, the running totals are divided by the total number of pixels that were processed (only those that survived filtering and are binarized to 1). This gives the average vegetation index calculation for that image, for each of the vegetation indices that were applied.

Eight vegetation indices (two of which had alternate forms) were applied, and a separate average was calculated for each of them for each of the plot-level images after filtering. The ExG has already been described, and was intended by its author to be used for partitioning crop images to distinguish soil from foliage^[6]^. As described previously, its alternate form tends to perform the best out of all the selected indices for distinguishing foliage from soil or shadows (**Figure 5**), and can be used to filter an image, but both of its forms can also be used as a greenness rating, just like each of the other indices. Other indices are the Red-Green-Blue Vegetation Index (RGBVI)^[3]^:

$$\frac{G^{2}-RB}{G^{2}+RB}$$

Which was intended to detect differences in absorption due to chlorophyll *a* and *b*^[3]^.

The Normalized Green-Red Difference Index (NGRDI)^[2]^:

$$\frac{G-R}{G+R}$$

Which has been used as a phenology indicator and has the potential for estimating biomass^[3]^, and also takes advantage of plants’ high reflectance in the green band, and low reflectance in the red and blue bands^[3,5]^.

The Modified NGRDI (MGRVI)^[3]^:

$$\frac{G^{2}-R^{2}}{G^{2}+R^{2}}$$

This index was developed from the NGRDI when it was observed that squaring the red and green reflectance values had the effect of amplifying the differences in reflectance between the red, green, and blue bands^[3]^.

The Green Leaf Index (GLI)^[4]^:

$$\frac{2G-R-B}{2G+R+B}$$

Which was intended to estimate the cover of wheat in digital images and works by determining if the average of the red and blue reflectance values is greater than or less than the green reflectance^[4]^. Essentially this is a partitioning index, like the ExG, and is meant to distinguish foliage from non-foliage elements.

The Visible Atmospherically Resistant Index (VARI)^[5]^:

$$\frac{G-R}{G+R-B}$$

This index was developed from the older NGRDI formula with an added correction for the blue reflectance. The authors state that this is intended to correct for atmospheric effects that may be present in satellite imagery. This index was also motivated by a perceived shortfall of the more widely-used NDVI, where near-infrared reflectance tends to level off in midseason and when the vegetation fraction (VF) reaches a certain point. As a result, the NDVI tends to not be sensitive to cases of high VF or LAI. The VARI was developed as a way to compensate for this problem by avoiding the use of near-infrared reflectance^[5]^.

The Vegetative Index (VEG)^[7]^:

$$\frac{g}{r^{a}b^{1-a}}$$

Where the variable *a* is said to depend on the center wavelengths of the camera’s three color filters. The original authors used a value of 0.667, and the same value was also used in this study^[7]^. The authors state that their intent with creating this index was not only to be able to partition images to distinguish soil from plants, but that the index would also be insensitive to varying light levels when the images were taken^[7]^.

Finally, the Woebbecke Index^[6]^ was used:

$$\frac{g-b}{|r-g|} , \frac{g-b}{r-g}$$

This index was introduced alongside the ExG as another potential equation to use for distinguishing foliage from soil^[6]^. The original paper did not name this index, but a paper by David and Ballado later referred to it as the Woebbecke Index^[8]^ after the original lead author, and so it is convenient to also refer to it as such here. This index has also been presented in two forms. Woebbecke presented the index with the absolute value in the denominator, while David and Ballado presented the equation without it. An earlier paper by Meyer and Neto also presents the index without the absolute value^[9]^.

**Supplementary Figures**

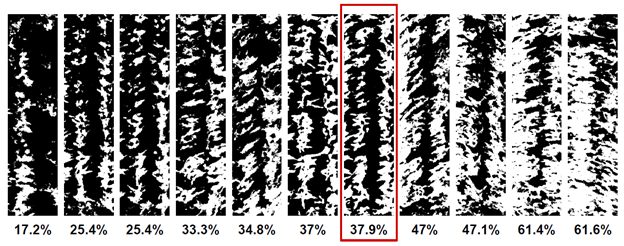

**Figure S1: Variability of plot replicate images.** These images represent all replicate images of an example plot after being filtered to remove non-foliage pixels. They have been sorted in increasing order by the percentage of white pixels, representing foliage. These percentages give an estimate of canopy coverage of the plot. However, depending on the viewing angle, this percentage can vary dramatically. It is important to use the reference replicate (red box) as a standard, as this guarantees better consistency when comparing different plots.

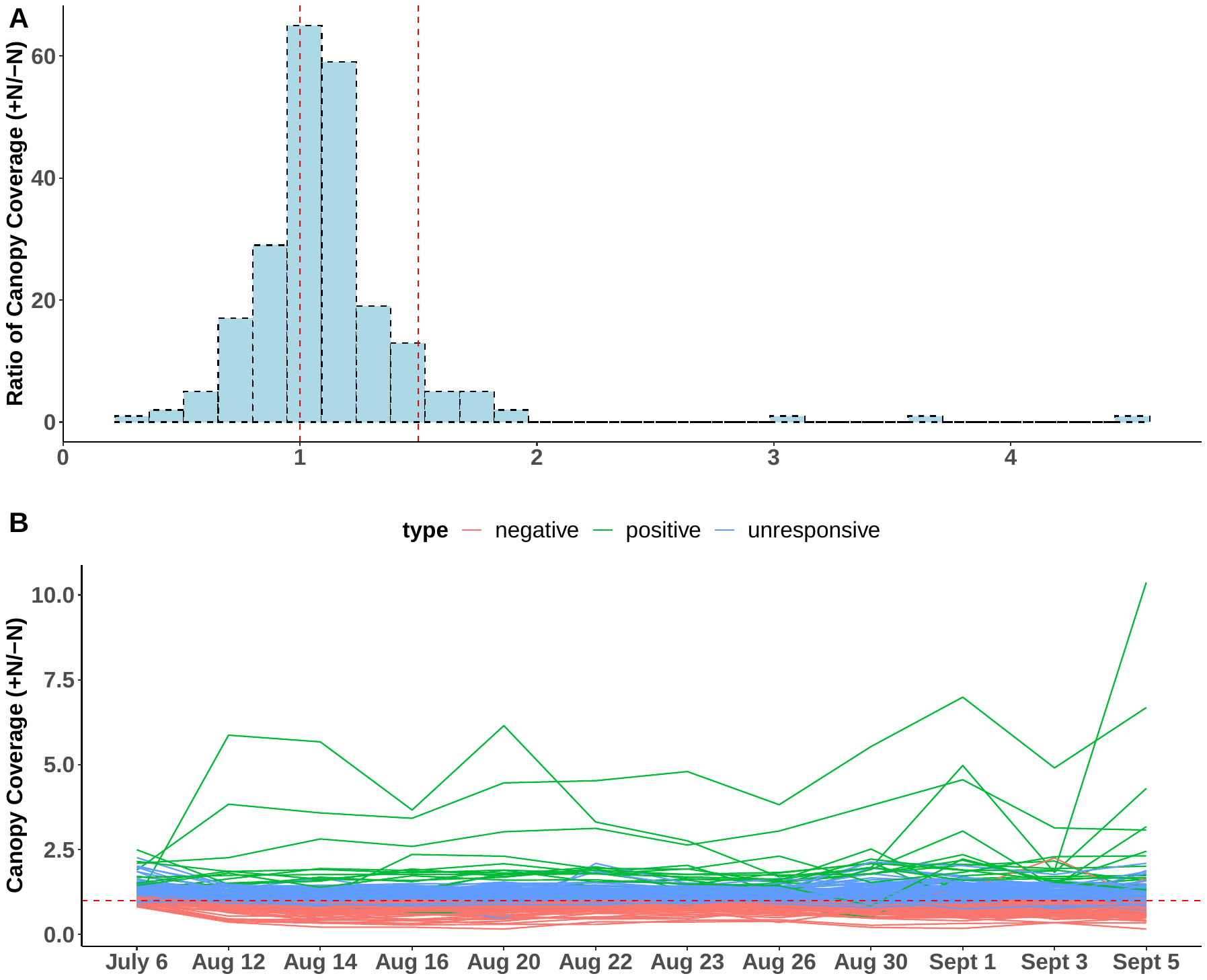

**Figure S2: Distributions of the ratios of canopy coverage for the 233 different genotypes.** (**A**) Histogram of the ratios of canopy coverage (+N/-N). Red dashed lines indicated the ratios of 1 and 1.5. (B) The time-series ratios of canopy coverage for genotypes with positive and negative responses to N treatments.

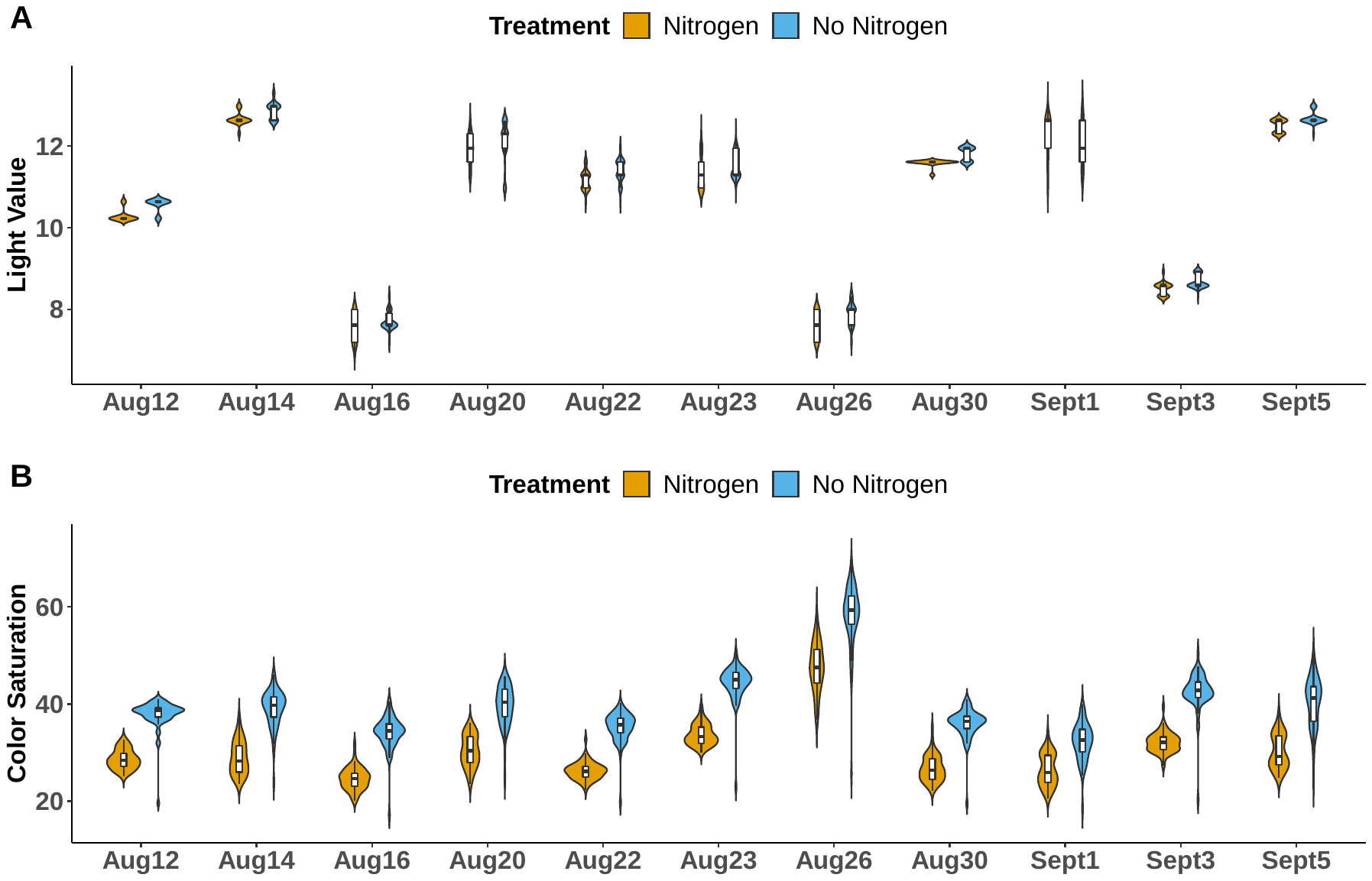

**Figure S3: Distributions of the light value and color saturation for the images of the hybrid check.** July 6 data was excluded from these plots because the values were fixed for that date.

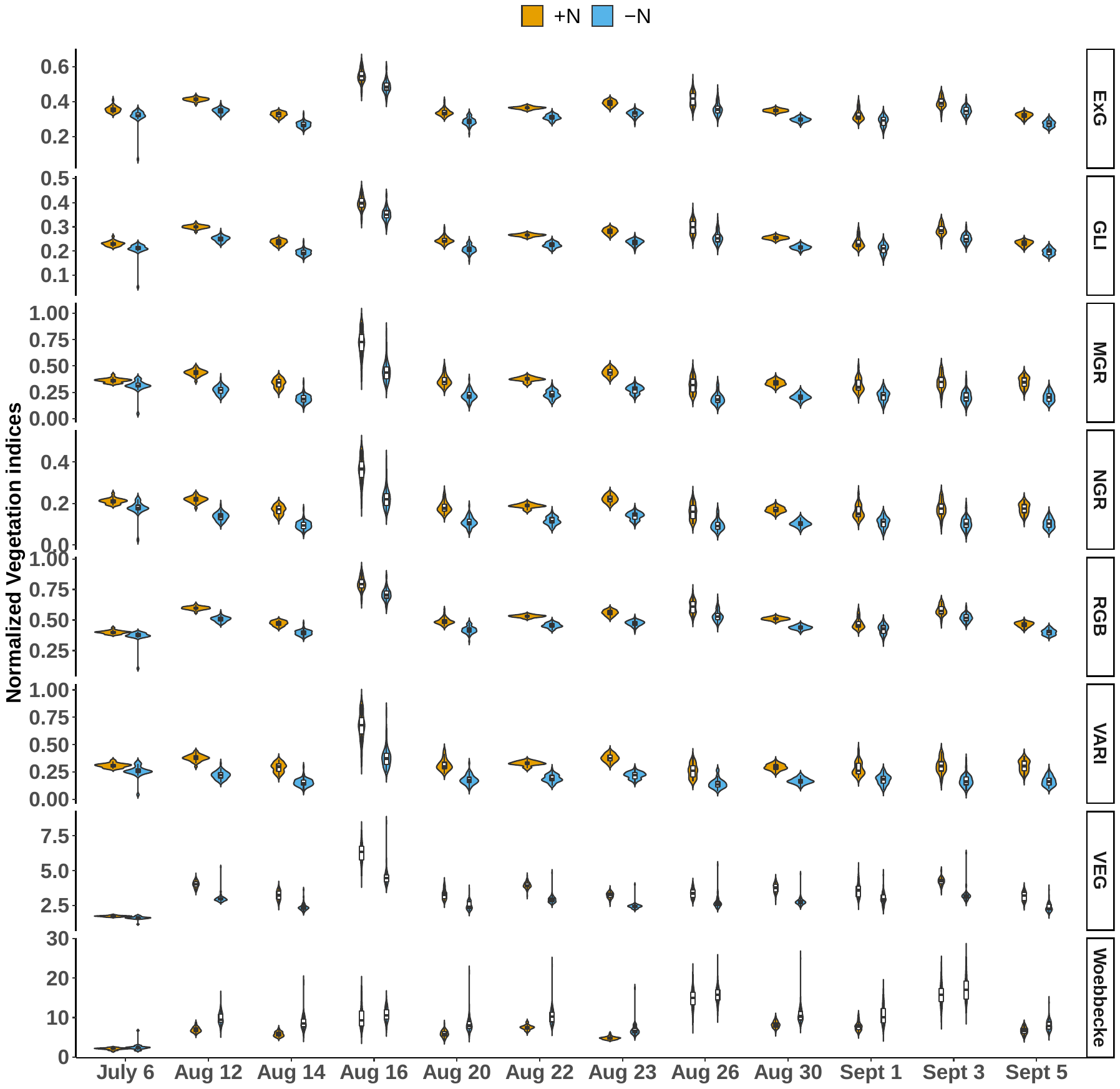

**Figure S4: Normalized vegetation indices of the hybrid checks under the +N and -N treatments.**

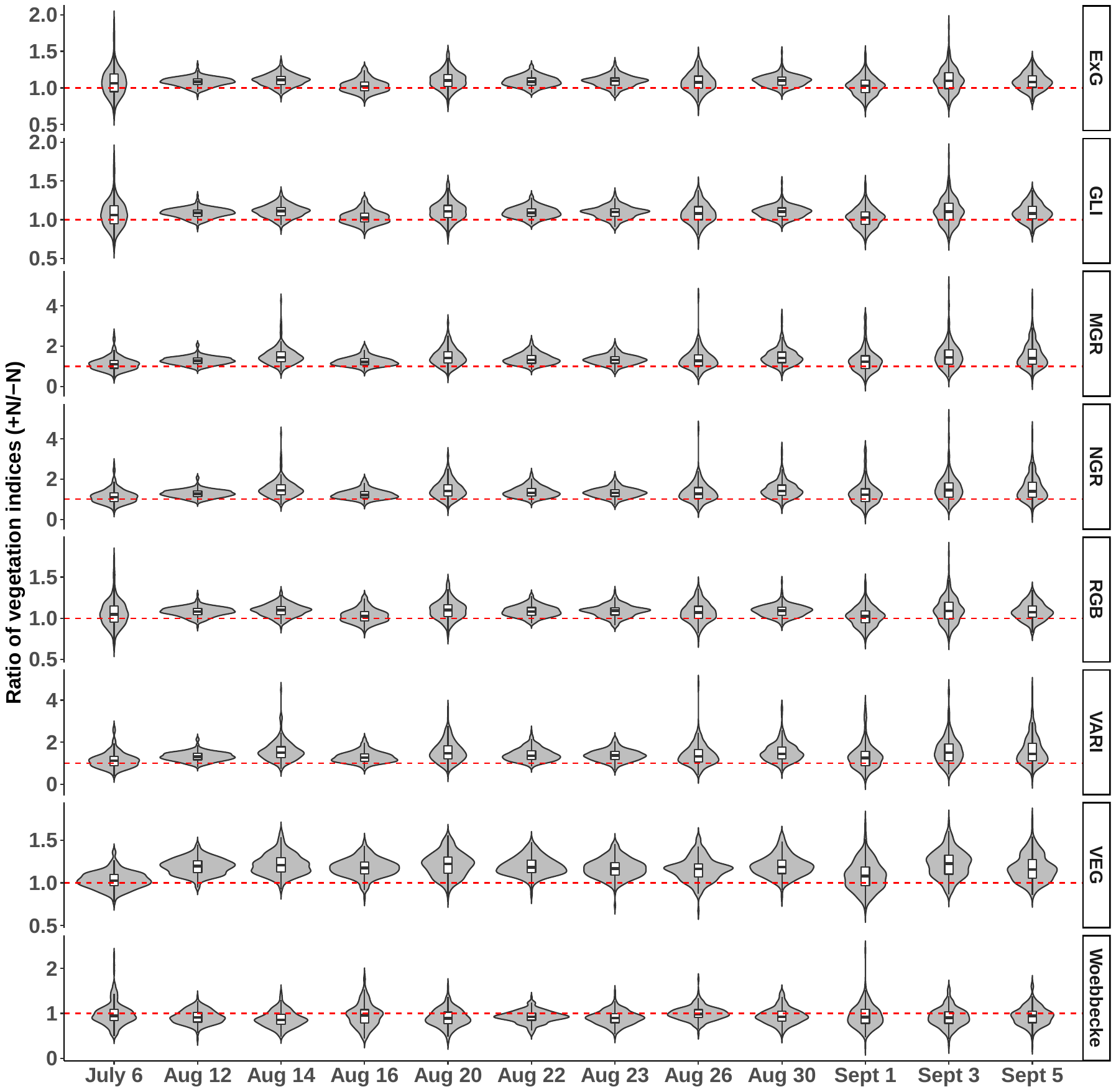

**Figure S5: Ratios of the normalized vegetation indices for the 233 genotypes.**

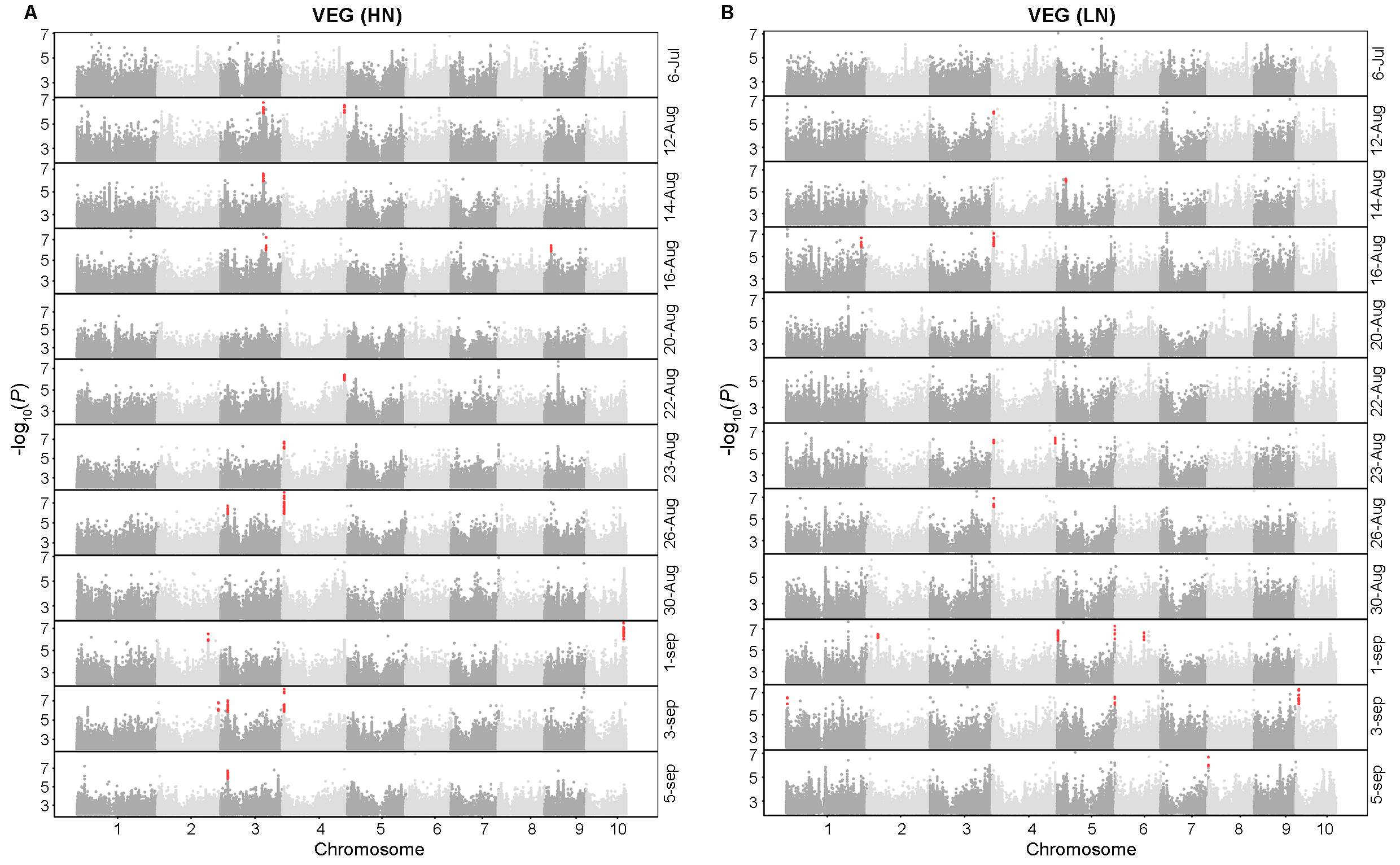

**Figure S6: GWAS results for Vegetative Index (VEG) at different dates with (A) and without nitrogen (B) treatments.** Red dots highlight the GWAS regions with more than five significant SNPs within a 100-kb window.
